# Synergistic Multi-Pronged Interactions Mediate the Effective Inhibition of Alpha-Synuclein Aggregation by the Chaperone HtrA1

**DOI:** 10.1101/2024.11.25.624572

**Authors:** Priscilla Chinchilla, Baifan Wang, Joseph H. Lubin, Xue Yang, Jonathan Roth, Sagar D. Khare, Jean Baum

**Author notes:** P.C. and B.W. contributed equally to this work.

## Abstract

The misfolding, aggregation, and the seeded spread of alpha synuclein (α-Syn) aggregates are linked to the pathogenesis of various neurodegenerative diseases, including Parkinson’s disease (PD). Understanding the mechanisms by which chaperone proteins prevent the production and seeding of α-Syn aggregates is crucial for developing effective therapeutic leads for tackling neurodegenerative diseases. We show that a catalytically inactive variant of the chaperone HtrA1 (HtrA1*) effectively inhibits both α-Syn monomer aggregation and templated fibril seeding, and demonstrate that this inhibition is mediated by synergistic interactions between its PDZ and Protease domains and α-Syn. Using biomolecular NMR, AFM and Rosetta-based computational analyses, we propose that the PDZ domain interacts with the C-terminal end of the monomer and the intrinsically disordered C-terminal domain of the α-Syn fibril. Furthermore, in agreement with sequence specificity calculations, the Protease domain cleaves in the aggregation-prone NAC domain at site T92/A93 in the monomer. Thus, through multi-pronged interactions and multi-site recognition of α-Syn, HtrA1* can effectively intervene at different stages along the α-Syn aggregation pathway, making it a robust inhibitor of α-Syn aggregation and templated seeding. Our studies illustrate, at high resolution, the crucial role of HtrA1 interactions with both the intrinsically disordered α-Syn monomers and with the dynamic flanking regions around the fibril core for inhibition of aggregation. This inhibition mechanism of the HtrA1 chaperone may provide a natural mechanistic blueprint for highly effective therapeutic agents against protein aggregation.

**Significance Statement:** PD and other synucleinopathies are marked by misfolding and aggregation of α-Syn, forming higher-order species that propagate aggregation in a prion-like manner. Understanding how chaperone proteins inhibit α-Syn aggregation and spread is essential for therapeutic development against neurodegeneration. Through an integrative approach of solution-based NMR, AFM, aggregation kinetics, and computational analysis, we reveal how a catalytically inactive variant of the chaperone HtrA1 effectively disrupts aggregation pathways. We find that the inactive Protease and PDZ domains of HtrA1 synergistically bind to key intrinsically disordered sites on both α-Syn monomers and fibrils, thereby effectively inhibiting both aggregation and templated seeding. Our work provides a natural and unique blueprint for designing inhibitors to prevent the formation and seeding of aggregates in neurodegenerative diseases.

## Introduction

Synucleinopathies, including Parkinson’s disease (PD), are debilitating age-related neurodegenerative diseases that affect millions of people worldwide (1). A pathological hallmark of PD and other synucleinopathies is neuronal cell death, which is associated with an abnormal accumulation of the intrinsically disordered protein (IDP) alpha synuclein (α-Syn) in amyloid-like fibrils (2, 3). Growing evidence suggests that α-Syn aggregates are transmitted between cells, leading to the templated growth of fibrils in a prion-like mechanism (4–6). α-Syn monomers are composed of three domains: an amphiphilic N-terminal domain (NTD), a hydrophobic NAC domain, and a highly-negatively charged C-terminal domain (CTD) (Fig. 1*A*) (4). In its fibrillar form, α-Syn adopts a “cross-beta” amyloid topology, with the NACore typically spanning residues 36-99, while the disordered NTD and CTD flank this core region (Fig 1*B*) (7–9). Insightful mechanistic studies of α-Syn aggregation inhibition have shown that the intrinsically disordered monomer, the fibril core, and the disordered flanking regions have all been targeted to disrupt aggregation and fibril formation (10–15). However, the intrinsic disorder of the α-Syn monomer and the disordered regions flanking the fibril core pose significant challenges for developing potent therapeutics. Several studies have explored proteins as inhibitors of α-Syn aggregation (16–20) and understanding the mechanisms by which natural proteins that are involved in cellular proteostasis inhibit aggregation may provide a blueprint for developing novel effective inhibitors to combat α-Syn-related pathologies.

**Figure 1:**
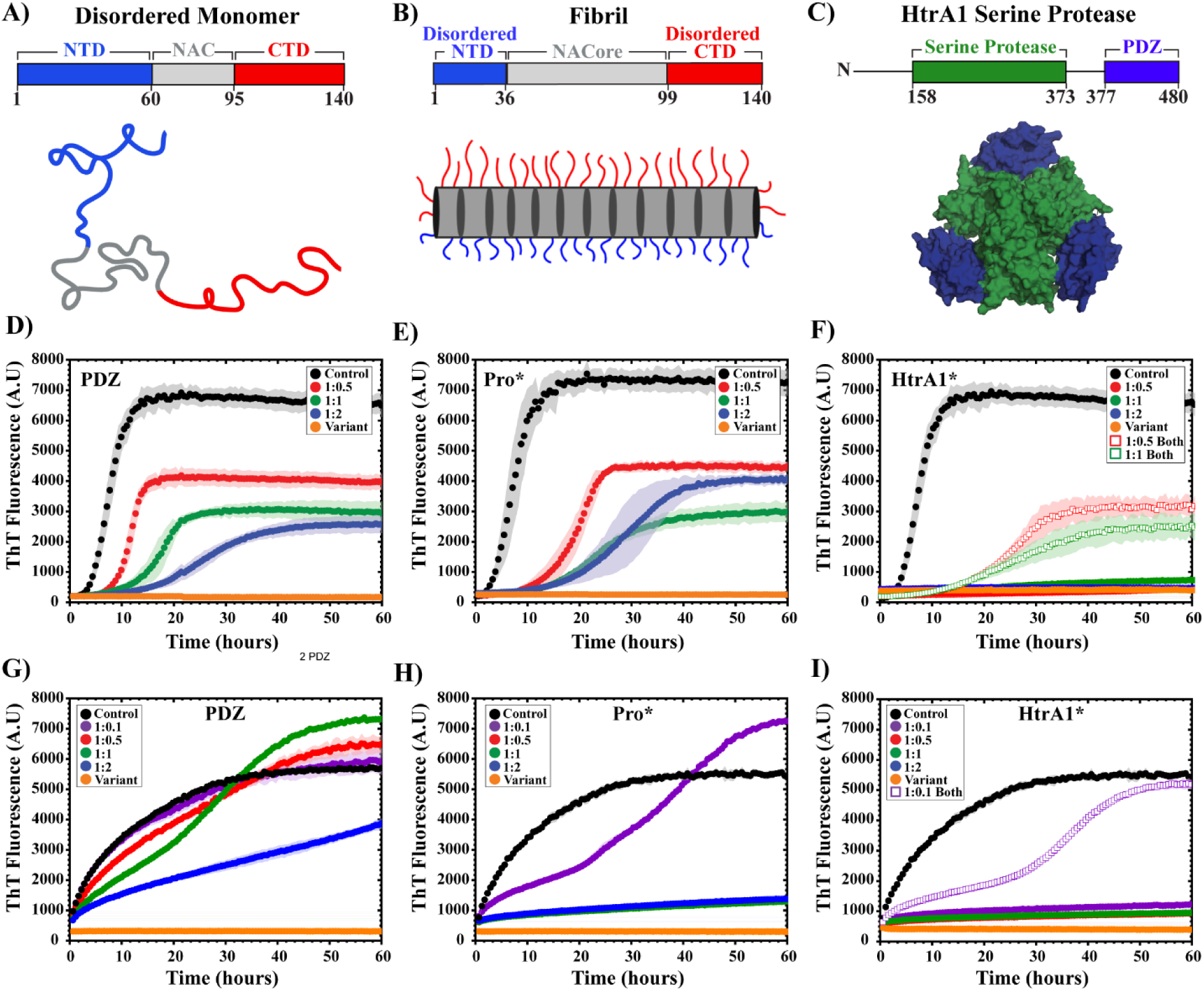
HtrA1* and its domain variants inhibit alpha synuclein (α-Syn) fibril formation *via de novo* and seeded aggregation assays. (**A**) Schematic of unbound, intrinsically disordered α-Syn, highlighting three domains: the amphipathic N-terminal domain (NTD, residues 1–60), the hydrophobic NAC region (residues 61–95), and the acidic C-terminal domain (CTD, residues 96– 140). (**B**) Schematic of α-Syn fibril with a disordered NTD (residues 1–36), a structured NACore (residues 37–99), and a disordered CTD (residues 100–140). In the fibrillar state, the NAC and part of the NTD form the core, while the remaining NTD and CTD remain disordered. (**C**) Trimeric HtrA1 model, highlighting the serine protease domain (green; residues 158–373) and C-terminal PDZ domains (blue; residues 377–480). The structure was generated by docking the protease domain (PDB: 3NZI) (62) with three PDZ domains (PDB: 2JOA) (35). (**D–F**) *De novo* fibril formation monitored by Thioflavin T (ThT) fluorescence. 50 µM α-Syn monomer was incubated with varying ratios of PDZ (**D**), Protease* (Pro*) (**E**), or HtrA1* (**F**) in 10 mM PBS (pH 7.4) at 37°C with constant agitation using a Teflon bead for 60 hours. Controls: α-Syn alone (black) and HtrA1 variants alone (orange). Ratios of α-Syn to each variant were 1:0.5 (red), 1:1 (green), and 1:2 (blue). For HtrA1* (**F**), empty red and green squares represent the addition of a combined 0.5 eq and 1 eq of PDZ and Pro*, respectively. (**G–I**) Seeded aggregation monitored by ThT fluorescence. 50 µM α-Syn monomer and 2.5 µM sonicated fibrils (seeds) were incubated with PDZ (**G**), Pro* (**H**), or HtrA1* (**I**) for 60 hours in 10 mM PBS (pH 7.4) at 37°C under quiescent conditions. Controls: α-Syn with seeds alone (black) and HtrA1 variants alone (orange). Ratios of α-Syn to each variant were 1:0.1 (purple), 1:0.5 (red), 1:1 (green), and 1:2 (blue). For HtrA1* (**I**), purple empty squares the addition of a combined 0.1 eq of PDZ and Pro*.

In humans, the high temperature requirement A (HtrA) family comprises three ATP-independent serine proteases (HtrA1-3) that have been shown to inhibit the aggregation of several amyloid-forming proteins including tau, amyloid beta (Aβ), and α-Syn (21–25). The HtrA family plays essential roles in various cellular processes including proliferation, cell migration, and cell maintenance (26, 27). Loss of HtrA activity has been linked to numerous diseases including cancer, arthritis, and neurodegenerative diseases (28–30). Among its family members, HtrA1 predominantly exists as a trimer composed of three domains: an N-terminal domain with IGFBP and Kazal-like modules, a serine Protease domain, and a C-terminal PDZ domain, which regulates protein-protein interactions (Fig. 1*C*) (31). These domains act on client proteins together, enabling a dual mode of action, in which HtrA1 functions both as a protease to degrade misfolded proteins and as a chaperone to fold substrates (27). In particular, HtrA1 has been shown to disaggregate and degrade different IDPs involved in the pathology of various neurodegenerative diseases (23–25). A recent comprehensive study revealed that HtrA1 inhibits the aggregation of α-Syn and remodels existing fibrils into non-toxic species that are incapable of seeding further aggregation (25). However, it is crucial to identify the molecular interactions through which HtrA1 and its domains engage with and disrupt α-Syn aggregation to illuminate the mechanisms of α-Syn aggregation inhibition and design targeted biologic therapeutics.

Here, using solution-based biomolecular Nuclear Magnetic Resonance (NMR) spectroscopy, computational modeling, Atomic Force Microscopy (AFM) and other biophysical approaches, we present the molecular mechanisms underlying inhibition of α-Syn aggregation by HtrA1 and delineate, at atomic resolution, the domain-domain interactions involved in these processes. Specifically, a catalytically inactive variant of HtrA1 (HtrA1*) effectively inhibits both the aggregation of α-Syn monomers and the seeding of fibrils. This inhibition is achieved through synergistic interactions between its PDZ domain, which targets the C-terminus of α-Syn, and its Protease domain, which targets the NAC region. In addition to the well-established role of the NAC domain, this work further highlights the critical gatekeeping function of the C-terminal domain in regulating α-Syn aggregation (32). Furthermore, we propose that HtrA1* inhibits fibril seeding through its interactions with the intrinsically disordered N- and C-terminal regions on the fibril surface, as well as by binding to α-Syn monomers. Thus, through multi-domain interactions and multi-site recognition of α-Syn, HtrA1* effectively intervenes at various stages of the aggregation pathway, functioning as a robust, multipronged inhibitor of both α-Syn aggregation and templated seeding. This work reveals a natural mechanistic framework for designing potent therapeutic agents that target the intrinsically disordered regions of α-Syn, offering a promising strategy to inhibit protein aggregation in synucleinopathies.

## Results

### HtrA1* and its individual domains inhibit *α*-Syn fibril formation as shown by Thioflavin-T fluorescence

Thioflavin-T (ThT) binding assays were performed to monitor the impact of HtrA1* and its domains on α-Syn fibril formation starting from purified monomeric protein. α-Syn fibrillation kinetics involve a lag phase that arises from a primary nucleation process, a rapid growth phase composed of secondary aggregation events, followed by a stationary phase (33). Inhibition of α-Syn aggregation by PDZ, an inactive Protease domain variant (Protease*), and HtrA1* exhibited significant dose dependence across all conditions (Fig. 1*D-F*). Both HtrA1* and its domain variants led to decreased aggregation rates; however, ThT fluorescence data reveal that PDZ alone is the least effective inhibitor, followed by Protease*. The two-domain construct HtrA1* demonstrates potent inhibition even at a 1:0.5 (α-Syn:HtrA1*) molar ratio. Notably, α-Syn incubated with HtrA1* shows significantly greater inhibition than the individual PDZ and Protease* domains added together (as separate proteins), suggesting that HtrA1* achieves enhanced efficacy through the proximity-induced synergy between its two domains.

The spread of pathological a-Syn aggregates likely occurs *via* templated seeding against pre-formed fibrils (PFFs) that are transmitted between cells (4–6). Therefore, we investigated the impact of HtrA1* and its domains on seeded aggregation (Fig. 1*G*-*I*). Seeded aggregation inhibition assays were performed to assess concentration dependent inhibition for the individual PDZ and Protease* domains, their combination, and HtrA1*. As with the unseeded aggregation kinetics (Fig. 1*D*-*F*), the inhibitory effects are dose-dependent and increased in the following order: PDZ, Protease*, the mixture of the two domains, and HtrA1* which was the most effective, achieving complete inhibition even at low concentrations (Fig. 1*G*-*I*). Notably, the PDZ domain alone does not efficiently inhibit seeded aggregation at substoichiometric concentrations (Fig. 1*G*). Consistent with the unseeded aggregation profiles, HtrA1* exhibited significantly greater inhibitory activity than PDZ and Protease* added together as separate proteins, suggesting that synergistic interactions between its two domains play a crucial role in inhibiting seeded aggregation as well.

### Tracking *α*-Syn aggregation with biophysical imaging confirms HtrA1*’s inhibition of fibril formation and growth

Although ThT fluorescence can monitor aggregation kinetics, it only detects higher-order aggregates with accessible binding pockets, leaving lower-order aggregates undetectable (34). To determine whether α-Syn treated with HtrA1* remains monomeric or forms ThT-incompetent aggregates, α-Syn monomers were incubated with an equimolar concentration of HtrA1* in 10 mM PBS (pH 7.4) at 37°C for one week followed by visualization using negative-stain Transmission Electron Microscopy (TEM) (Fig. 2*A*). Incubation of α-Syn alone resulted in mature fibril formation, whereas HtrA1* formed amorphous structures with quasi-spherical morphology. Interestingly, α-Syn monomers treated with HtrA1* produced insoluble amorphous aggregates, appearing to be surrounded by HtrA1*. Similar aggregates produced by HtrA1* treatment have previously been shown to be non-toxic and incapable of seeding aggregation when transfected into primary mouse neurons (25).

**Figure 2:**
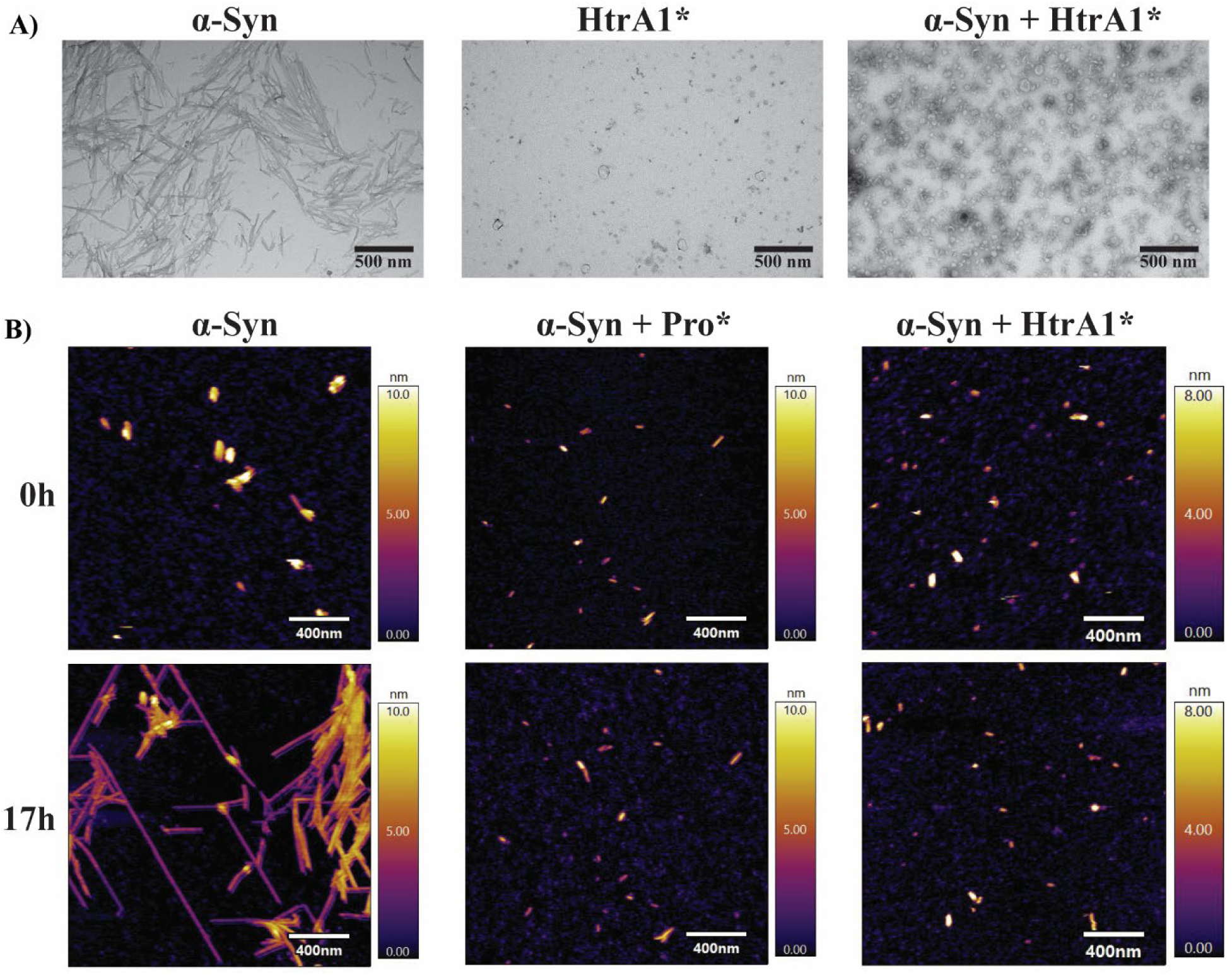
Tracking α-Syn aggregation and its modulation by HtrA1 through biophysical imaging techniques. (**A**) Negatively stained TEM images of 50 μM α-Syn monomer, 50 μM HtrA1*, and a mixture of 50 μM α-Syn and HtrA1* after one week of incubation in 10 mM PBS (pH 7.4) at 37°C, 900 rpm using a Thermomixer C (Eppendorf). Incubation of α-Syn alone resulted in mature fibril formation, while HtrA1* produced small spherical structures. Co-incubation of α-Syn with HtrA1* yielded insoluble amorphous aggregates. Images were captured using a JEOL JEM2010 TEM at 22,000× magnification with a 500 nm scale bar. (**B**) Liquid AFM time-lapse images taken at 0 and 17 hours, tracking seeded α-Syn fibril growth in the absence and presence of Protease* (Pro*) and HtrA1*. At timepoint 0, the presence of 5 μM sonicated fibrils (seeds) was confirmed using AFM imaging. Following confirmation, a solution of 25 μM α-Syn monomer in the absence and presence of 1 μM of HtrA1 variants was added to the seeds in 10 mM PBS (pH 7.4), at 25°C. In the control condition (α-Syn monomers and seeds), rapid fibril growth was observed over 17 hours. The addition of Pro* partially inhibited fibril growth, while HtrA1* appeared to prevent fibril formation entirely, with no visible growth detected. All AFM imaging was performed using a Cypher ES AFM (Asylum Research, Oxford Instruments) in tapping mode with blueDrive photothermal excitation for enhanced liquid imaging resolution.

To further investigate how HtrA1 variants affect α-Syn aggregation, AFM was used to monitor their impact on α-Syn seeded aggregation in real-time. For these experiments, 25 μM α-Syn monomers and 5.0 μM sonicated fibrils (seeds) were incubated with and without 1 μM Protease* and HtrA1* in 10 mM PBS (pH 7.4) at 25°C and imaged continuously for 17 hours using a Cypher ES AFM in liquid mode (Fig. 2*B*). Following imaging, individual AFM frames were stitched together to create videos illustrating the aggregation dynamics, which are available in the SI Appendix as movie files (Movies S1–S3). In the α-Syn-only control, elongation and secondary nucleation resulted in the formation of mature fibrils (Fig. 2*B*, Movie. S1). Notably, areas that were continuously imaged by the AFM tip grew significantly faster than areas that were not imaged. This suggests that the tip may be causing nano-fractures in the incipient seeds and fibril structures, thereby enhancing seeded aggregation. Despite being imaged under the same conditions, α-Syn incubated with Protease* showed a few elongation events but did not form mature fibrils (Fig. 2*B*, Movie. S2). Strikingly, α-Syn incubated with HtrA1* showed no significant aggregation (Fig. 2*B*, Movie. S3). These findings demonstrate that even at very low stoichiometries (25:1 monomer:HtrA1* and 5:1 seeds:HtrA1*) and in the presence of aggregation-promoting mechanical forces, HtrA1* effectively inhibits seeded aggregation. Together, AFM, TEM, and fibrillation kinetics confirm that HtrA1* and its domains disrupt α-Syn fibril formation and growth. Furthermore, this study highlights the use of AFM as a powerful tool for observing and analyzing the formation of higher-order protein structures, such as α-Syn fibrils, in real time.

### HtrA1 engages with the intrinsically disordered C-terminal domain of *α*-Syn monomers through its PDZ domain

To investigate the residue-level interactions underlying HtrA1*’s inhibition of α-Syn aggregation, interactions α-Syn interactions with HtrA1 variants were characterized. Using solution-based NMR, residue-specific interactions between α-Syn monomers and the PDZ domain were examined, as PDZ domains are known to recognize C-terminal residues of IDPs and mediate HtrA1’s role in protein-protein interactions (35–37). This approach enabled us to assess PDZ’s role in selectively binding to α-Syn regions involved in aggregation. NMR transverse relaxation experiments were conducted on ^15^N labeled α-Syn monomers incubated with 1 equivalent (eq) of unlabeled PDZ (Fig. 3*A*). A comparison of ^15^N R_2_ data showed complete peak broadening for residues 136-140 of α-Syn, indicative of specific binding at these sites. The PDZ domain of HtrA1 predominately recognizes hydrophobic residues at the C-terminus of protein chains (35–37). This is consistent with our findings that PDZ interacts with the last four C-terminal residues of α-Syn, two of which are hydrophobic (Alanine and Proline) (Fig. 3*A*). Furthermore, the addition of PDZ showed extended line broadening and elevated ^15^N R_2_ values within α-Syn’s C-terminal residues 110-140 indicating that PDZ engages in weak, broad interactions with the C-terminal domain of α-Syn, a region critical for modulating aggregation (10, 32, 38–40).

**Figure 3:**
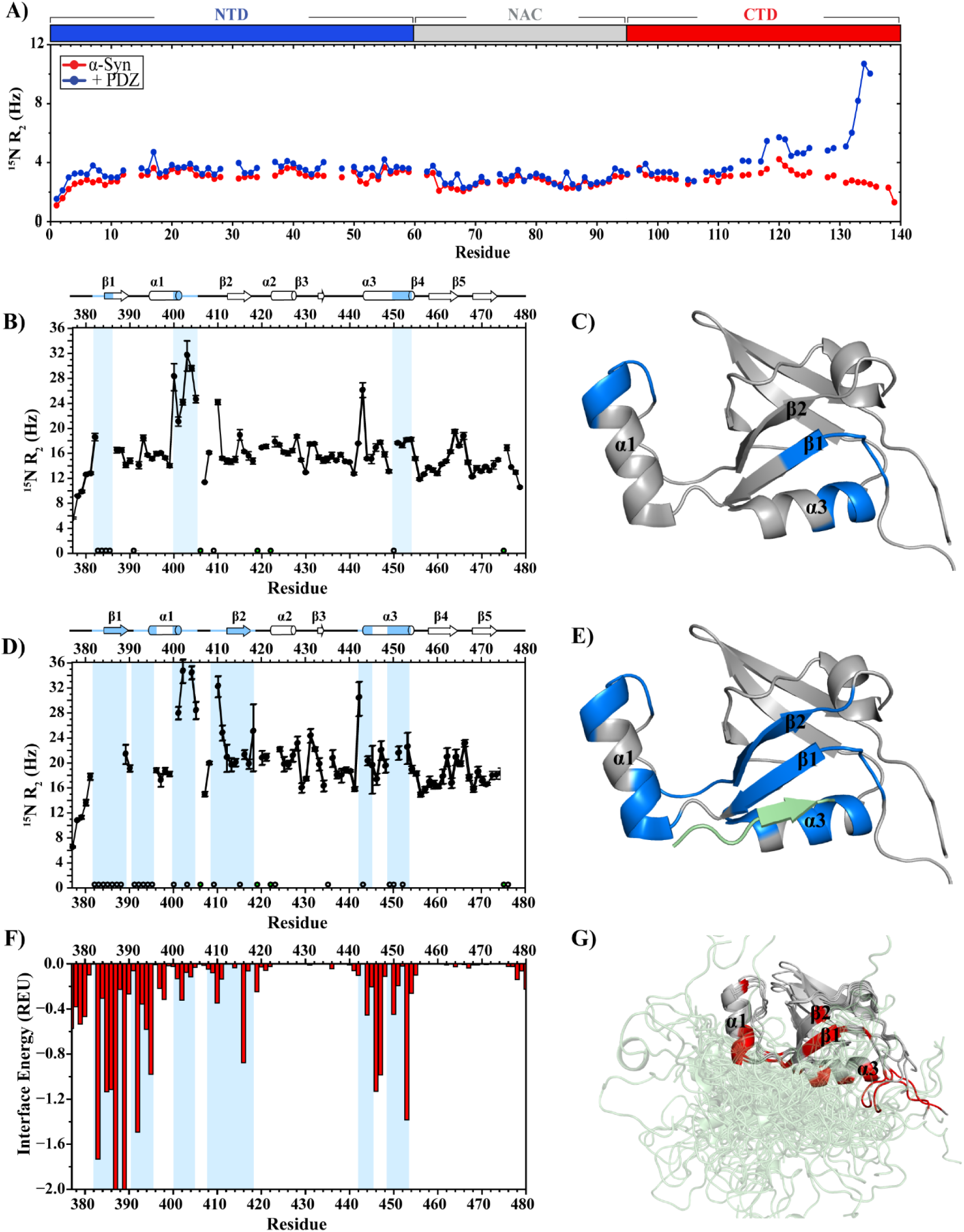
The PDZ domain of HtrA1 interacts with the intrinsically disordered C-terminus of α-Syn monomers. (**A)** ^15^N R_2_ solution NMR data for α-Syn monomers in the absence (red) and presence of 1 eq of PDZ (blue). 200 μM of ^15^N-labeled α-Syn monomers and 200 μM of unlabeled PDZ were mixed in NMR buffer (20 mM MES, 100 mM NaCl, pH 6.0, 10% D2O). Spectra were acquired on an 800 MHz Varian Inova spectrometer. Elevated ^15^N R_2_ values and broadening at the α-Syn C-terminus in the presence of PDZ indicate potential interaction sites. (**B, D**) ^15^N R_2_ solution-based NMR data for ^15^N-labeled PDZ in the free state (**B**) and in the presence of 1 eq unlabeled α-Syn monomer (**D**). 200 μM of ^15^N-labeled PDZ and 200 μM of unlabeled α-Syn monomer were mixed in NMR buffer and spectra were acquired on an 800 MHz Varian Inova spectrometer. Residues with elevated ^15^N R_2_ values or unobservable signals (three or more consecutive) are highlighted in blue on the spectra. These regions are mapped onto PDZ’s secondary structures shown above the spectra. Elevated ^15^N R_2_ values are defined as those exceeding 1 standard deviation above the mean, with outliers removed using the interquartile method. Significant broadening and elevated ^15^N R_2_ values were observed in α1, α3, β1, and β2 regions, suggesting interactions with α-Syn at the PDZ binding pocket and its neighboring structures. Open circles denote unassigned/unobservable residues, and green circles indicate proline residues. (**C, E**) Structural mapping of PDZ (PDB: 2JOA) (35) highlighting areas corresponding to elevated ^15^N R_2_ values in the free state (**C**) and the α-Syn-bound state (**E**). The bound peptide (DSRIWWV), shown in green, models α-Syn’s C-terminal interactions with PDZ. (**F**) Interface energy spectra of PDZ residues bound to α-Syn monomers, calculated using Rosetta modeling. Blue shading highlights regions with elevated ^15^N R_2_ values from the α-Syn-bound PDZ spectrum (**C**) and corresponds well to residues with low interface energies, indicating their involvement in PDZ-α-Syn interactions. (**G**) Conformational ensemble of the 100 lowest-energy models of α-Syn residues 110-140 (green) bound to PDZ. Red highlights indicate residues with interface energies lower than 1 standard deviation below the mean (excluding outliers). These residues localize primarily to α1, α3, β1, and β2, supporting the observed NMR data and indicating their involvement in binding.

### NMR and computational modeling reveal weak, dynamic interactions between PDZ and the disordered C-terminal region of *α*-Syn monomers

After observing that PDZ interacts with the C-terminal domain of α-Syn, transverse relaxation experiments with ^15^N-labeled PDZ and unlabeled α-Syn monomers were conducted to delineate the PDZ residues involved in α-Syn binding. In a previously determined peptide-bound structure of the HtrA1 PDZ domain (PDB: 2JOA), hydrophobic residues in β1 and α3 helix regions mediate direct contacts with the peptide (35). For the unbound PDZ (Fig. 3*B*,*C*), highly elevated ^15^N R_2_ rates are present for residues at the C-terminal end of the α1 helix into the loop connecting α1 with β2, as well as elevated values in α3, and missing resonances (residues 383-386) in the N-terminus leading to β1, potentially due to conformational exchange. Addition of α-Syn monomers further broadens resonances at residues 382-388 and 391-395 around β1 and the α1 N-terminus, with increased broadening between α1 and β2; elevated ^15^N R_2_ rates are retained in the α3 helix suggesting that α-Syn interacts in the PDZ binding pocket encompassing α3 and β1 (Fig. 3*D*,*E*). Additionally, given its intrinsically disordered nature, the α-Syn C-terminal domain likely transiently interacts with residues in neighboring structures near PDZ’s binding pocket, including those in α1 and β2.

Computational modeling was used to analyze α-Syn binding to PDZ, calculating binding energy contributions from PDZ residues using Rosetta (Fig. 3*F*,*G*). This approach allowed us to capture the distinct energetic contributions involved in IDP binding, which differ from those in traditional interactions with fully folded proteins (41, 42). Favorable binding energies were observed for PDZ residues in the β1, the N-terminus of α1, β2, and α3 (Fig. 3*G*). Comparison of elevated ^15^N R_2_ values with regions showing favorable binding energies (Fig. *3F*) suggests that direct interactions between PDZ and α-Syn may underlie the elevated NMR ^15^N R_2_ values. Through NMR relaxation experiments and computational modeling, PDZ binding to α-Syn monomers is revealed to be highly dynamic, involving flexibility changes in specific regions that extend beyond the peptide-binding site observed in the PDZ domain crystal structure (Fig. 3*G*) (35).

### HtrA1 cleaves the NAC domain of *α*-Syn monomers with the first cleavage occurring between residues T92 and G93

Previous studies and our work have shown that the proteolytically inactive version of Protease (Protease*) can delay α-Syn fibril formation (Fig. 1*E*,*H*) (25), indicating a direct interaction between Protease and α-Syn. To identify the specific interactions between α-Syn and Protease, we utilized the catalytically active HtrA1, where α-Syn monomers were incubated with HtrA1 at a 5:1 ratio in 50 mM Tris-HCl (pH 8.0) at 37°C, followed by SDS-PAGE analysis (Fig. 4*A*). Within minutes, band intensity for α-Syn diminished, and new bands corresponding to proteolytic fragments appeared around 9-10 kDa. Mass spectrometry (MS) identified the proteolytic products of α-Syn (Fig. 4*B*). Specifically, α-Syn treated with HtrA1 at a 5:1 ratio for 10 minutes exhibited a prominent mass peak of 9130.5 Daltons, corresponding to residues M1-T92 of α-Syn (Table 1). Further incubation for four hours revealed additional fragments, showing that HtrA1 continues cleaving beyond the initial cut.

**Figure 4:**
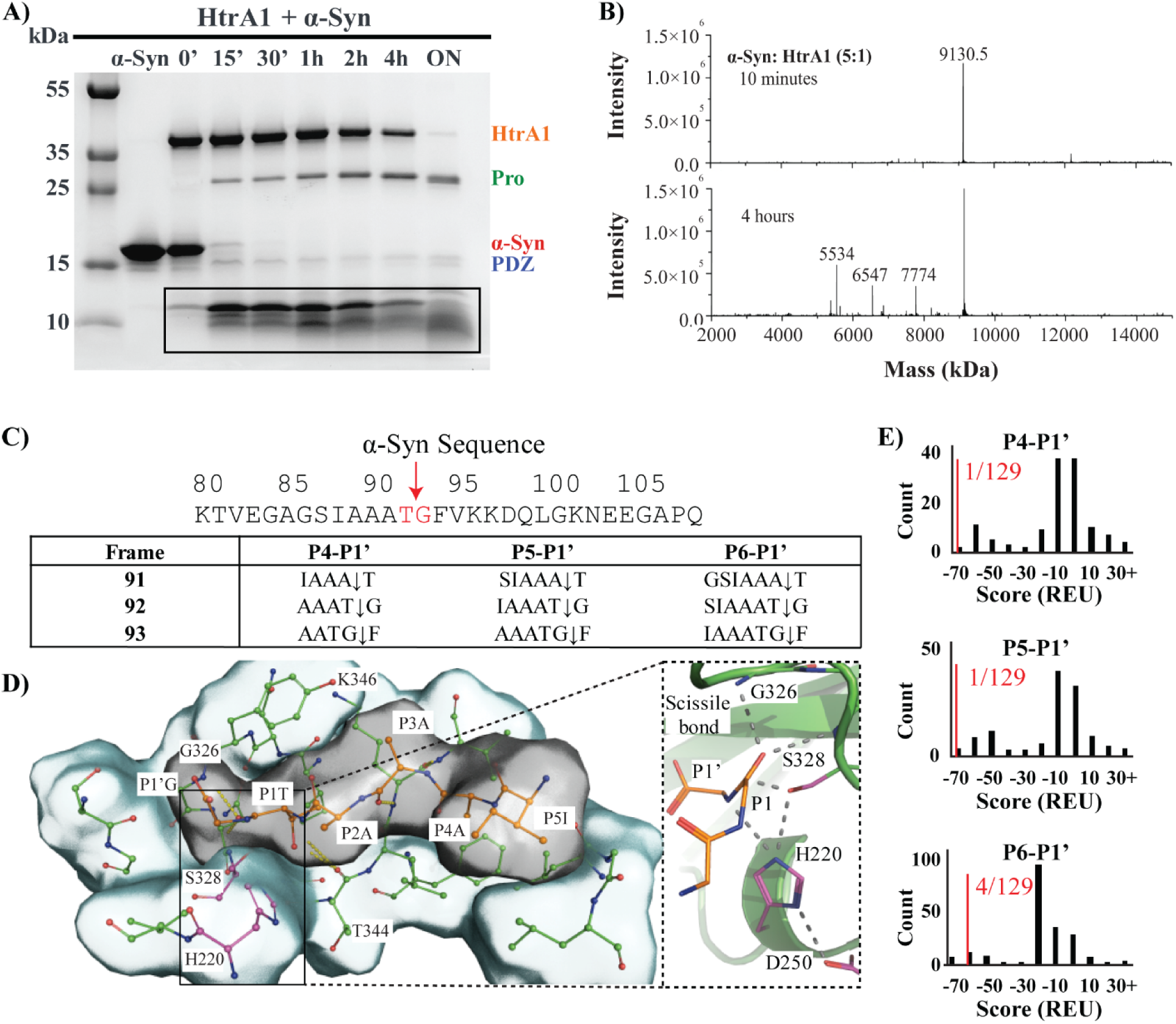
HtrA1 cleaves α-Syn monomers with specificity between residues T92 and G93. (**A**) Proteolysis of α-Syn by HtrA1 visualized using SDS-PAGE. α-Syn monomer was incubated with HtrA1 at a 5:1 ratio across various timepoints in 50 mM Tris-HCl (pH 8.0) at 37°C. Cleavage fragments of α-Syn were detected at 0’, with most of the monomer cleaved after overnight incubation. Bands corresponding to HtrA1 (orange), Pro (green), PDZ (blue), and α-Syn (red) are labeled accordingly. Cleaved α-Syn fragments are highlighted with a black box. (**B**) Mass spectrometry identification of HtrA1-mediated α-Syn cleavage. α-Syn monomer was incubated with HtrA1 at a 5:1 ratio in 50 mM Tris-HCl (pH 8.0) at 37°C for 10 and 4 hours, yielding a main peak at 9130.5 Da, corresponding to the 1–92 α-Syn fragment, which confirms cleavage specificity. Prolonged incubation resulted in additional peaks, suggesting further cleavage events following the initial cleavage between residues T92 and G93. (**C-E**) Computational modeling of Protease-bound α-Syn. (**C**) Frames of the α-Syn sequence were modeled in catalytic geometry with varying peptide lengths. The smallest modeled peptide, P4-P1’, included four residues upstream of the scissile bond and one downstream for a total of five residues. All 5-residue frames of the α-Syn sequence were modeled, with the sequence at site 92 being ‘AAATG’. (**D**) P5-P1’ of the cleaved site 92 (sequence IAAATG) in the active site of the Protease. The catalytic geometry of the Protease was enforced by constraints between all catalytic residues, the oxyanion hole, and the P1 and P1’ residues of the peptide. (**E**) Discriminator histograms for three representative peptide sizes. Histograms represent the discriminator scores of all frames in the α-Syn sequence. The red line and text indicate the discriminator value for site 92, consistent with experimental cleavage results.

**Table 1:** Fragments of α-Syn identified through MS, detailing fragment lengths, expected and observed masses, delta values representing mass differences, and corresponding cleavage sites.

| Identified Fragments | $[M]_{\text{exp.}}$ | $[M]_{\text{calc.}}$ | $\Delta[M]$ | Cleavage Sites |
| --- | --- | --- | --- | --- |
| 1-92 | 9130.5 | 9130.48 | -0.017 | T(G) |
| 1-77 | 7774.0 | 7773.98 | -0.016 | V(A) |
| 1-63 | 6547.5 | 6547.59 | 0.087 | V(T) |
| 1-54 | 5534.5 | 5534.43 | -0.071 | T(V) |
| 1-140 | 14502.5 | 14509.19 | -0.306 | n.a. |

In a complementary analysis, an HSQC conducted on ^15^N labeled α-Syn monomers treated with 1 eq of unlabeled Protease showed significant chemical shift perturbations in residue T92, while residue G93 became unobservable (Fig. S1*A*), suggesting that the Protease domain cleaves the peptide bond between T92 and G93. To compare the proteolytic activity of HtrA1 and Protease on α-Syn, similar SDS-PAGE and MS analyses were performed for Protease treatment (Fig. S1*B*,*C*). Results showed similar proteolytic properties for HtrA1 and Protease on α-Syn. However, while overnight incubation with HtrA1 and Protease showed comparable cleavage, kinetic analyses revealed that HtrA1 cleaves significantly faster than Protease, due to synergistic contributions from the PDZ domain, which may anchor α-Syn and increase accessibility to HtrA1’s initial cleavage site. It has been found that cleaving C-terminal domain of α-Syn monomer results in facile aggregation of the N-terminal fragment, if the cleavage occurs after residue 99 (43–45). We investigated whether the products of proteolytic cleavage have a propensity to aggregate and found that the protein fragment consisting of residues 1-92 is incapable of aggregating on its own (Fig. S1*D*).

To investigate the molecular basis for the cleavage site preference at residue 92, Rosetta-based calculations of protease-substrate complexes were performed. Overlapping fragments of α-Syn were generated from scanning its sequence with different variable-length windows and each peptide fragment was modeled to bind within the Protease active site of HtrA1 (Fig. 4*C*) using a protocol we previously developed for predicting protease specificity (46). A protease specificity discriminator (PSD) score was calculated for each of the α-Syn fragments bound to HtrA1 (Fig. 4*D*). The cleavage sites of each α-Syn fragment were ranked by the single best discriminator value determined among the ten lowest-energy structures (Fig. 4*E*). The PSD score identified residue 92, the experimentally observed first cleavage site, as the most energetically favorable frame in the α-Syn sequence. Taken together, our results suggest that the ability of the Protease to bind in a catalytically productive conformation underlies the observed specificity for that site and the presence of the PDZ domain serves to enhance the accessibility of the first cleavage site.

### PDZ binds selectively to the fuzzy coat of the C-terminal domain of *α*-Syn fibrils

α-Syn aggregation inhibition may arise *via* interactions between HtrA1 and either the monomer, the fibril, or both. Following the identification of PDZ binding sites on α-Syn monomers, the interactions between PDZ and α-Syn fibrils were investigated. A pulldown experiment demonstrated dose-dependent binding of PDZ to α-Syn fibrils (Fig. 5*A*). Next, NMR experiments were conducted to investigate interactions between PDZ and α-Syn fibrils at the residue level (Fig. 5*B-D*). For these experiments, ^15^N labeled PDZ was titrated with 0.25 and 0.35 eq of unlabeled α-Syn fibrils with HSQC spectra revealing dose-dependent peak broadening across all PDZ residues likely resulting from slower tumbling upon PDZ binding to the large α-Syn fibrillar species (Fig. 5*B)*. NMR transverse relaxation experiments were used to further pinpoint specific PDZ residues that are affected by increasing fibril concentrations (Fig. 5*C*,*D*). Addition of 0.25 eq of fibrils caused elevated ^15^N R_2_ values in residues of β1, β2, and the C-terminus of β3 in PDZ. Upon the addition of 0.35 eq of fibrils, some residues in β1and β2 structures that previously had elevated ^15^N R_2_ values became unobservable (Fig. 5*D*). Additionally, new peaks corresponding to residues in β1, α2 and β3 structures broadened as well. This broadening and elevated ^15^N R_2_ values in PDZ’s β1, β2, and β3 regions upon fibril titration suggest dose-dependent binding to the C-terminal “fuzzy coat” of α-Syn fibrils (Fig. 5*C*,*D*) resembling interactions observed with α-Syn monomers (Fig. 3*D*,*E*). The elevated ^15^N R_2_ values and peak broadening observed in α2 and β3 residues suggest that the PDZ front surface may also participate in binding with α-Syn fibrils (Fig. 5*E*).

**Figure 5:**
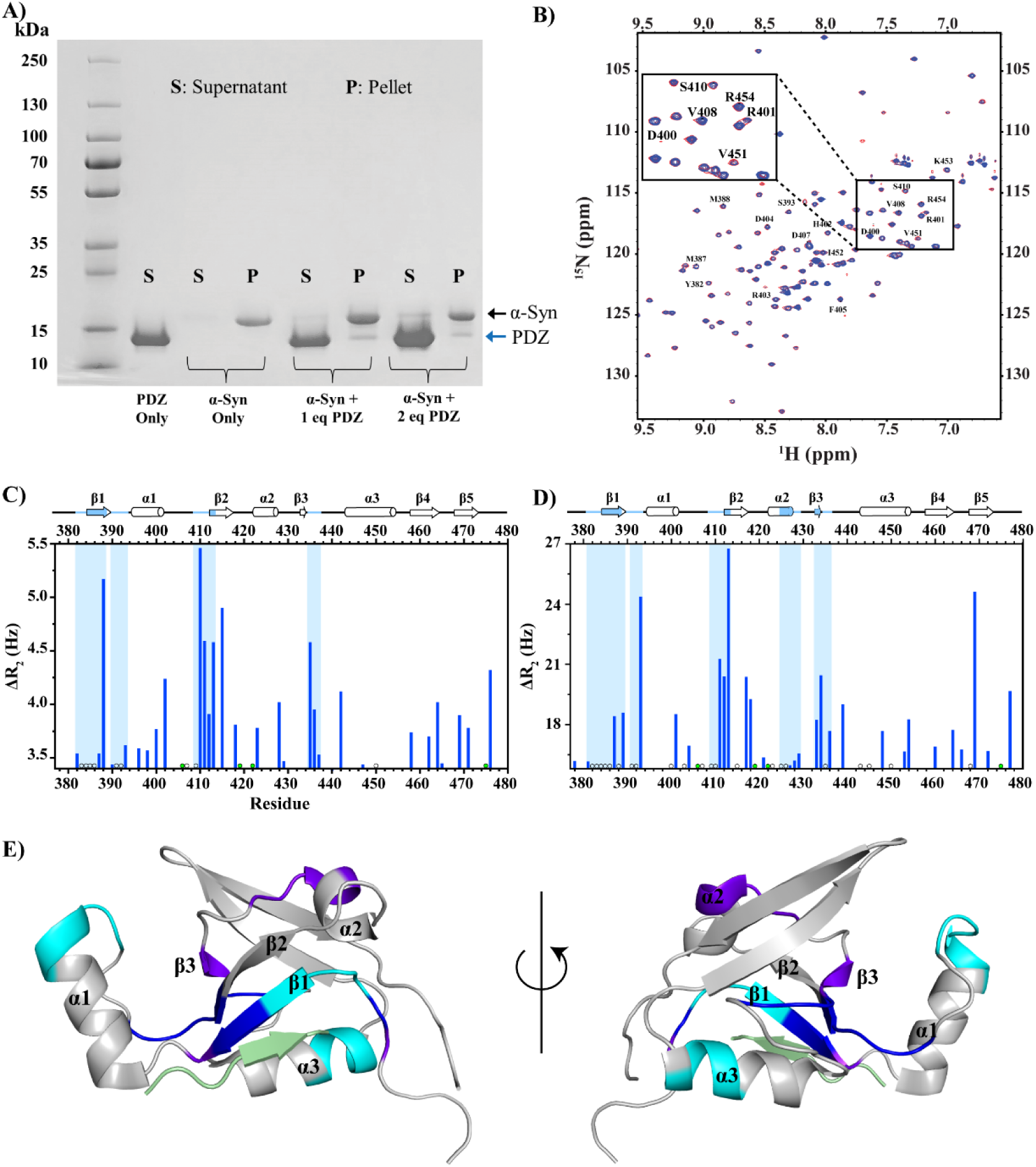
PDZ transiently interacts with the fuzzy coat of α-syn fibrils. (**A**) PDZ pulldown assay demonstrating the complex formation between PDZ and α-Syn fibrils. 50 or 100 μM PDZ was incubated with 50 μM α-Syn fibrils in 10 mM PBS (pH 7.4) at 37°C for 24 hours with shaking. Control samples containing only 50 μM PDZ or 50 μM α-Syn fibrils were incubated under identical conditions. After incubation, the PDZ-fibril complex was isolated by centrifugation. The supernatant (S) contained unbound proteins, while the pellet (P) contained any protein complexes formed. Pelleted samples were denatured overnight and analyzed by SDS-PAGE alongside the supernatant samples. The presence of α-Syn and PDZ in the pellet, as shown by SDS-PAGE, confirms the formation of a PDZ-fibril complex. Bands corresponding to α-Syn and PDZ are labeled. (**B**) ^15^N-^1^H HSQC spectra of 200 μM ^15^N labeled PDZ in the absence (red) and presence of unlabeled 0.25 eq α-Syn fibrils (blue). A magnified view highlighting peaks corresponding to key residues within PDZ’s β2, α1, and α3 regions is shown in the top corner. (**C**, **D**) NMR transverse relaxation experiments were performed with 200 μM PDZ titrated with 0.25 and 0.35 eq unlabeled fibril concentrations and Δ^15^N R_2_ values of ^15^N-labeled PDZ were calculated as the difference between fibril-bound and unbound PDZ. The results were plotted above a baseline, defined as 0.25 SD above the mean, with outliers removed using the interquartile method. Regions with three or more consecutive residues showing elevated Δ^15^N-R_2_ values above the baseline, or with unobservable signals, are highlighted in blue on the spectra and mapped onto the corresponding PDZ secondary structures displayed above the spectra. Significant broadening and elevated Δ^15^N R2 values were detected in the β1 and β2 regions, indicating interactions between α-Syn fibrils and the PDZ binding pocket. Similar values in the α2 and β2 regions, located on the front of PDZ, further support that fibrils engage with both the binding pocket and the front surface of the PDZ domain. Open circles denote unassigned/unobservable residues, and green circles indicate proline residues. All NMR data was acquired in 20 mM, 100 mM NaCl (pH 6) with 10% D2O and 15 °C at 700 MHz ^1^H Larmor frequency. (**E**) The structural mapping of PDZ (PDB: 2JOA) (35) highlights regions with elevated Δ^15^N-R_2_ values: cyan for the free state, dark blue for the 0.25 eq fibril-bound state (**C**), and purple for the 0.35 eq state (**D**). The bound peptide, DSRIWWV (green), models the binding site for α-Syn fibrils. Highlighted areas indicate that α-Syn fibrils primarily interact with the front surface of PDZ, as modeled by the bound peptide (DSRIWWV, green).

### HtrA1 binds the flanking N- and C-terminal domains of *α*-Syn fibrils, potentially modulating aggregation

Following confirmation of PDZ binding to α-Syn fibrils, NMR was used to map regions of α-Syn fibrils interacting with HtrA1. For this experiment, ^15^N labeled α-Syn fibrils were treated with 2.5 eq of HtrA1 in 50 mM Tris-HCl (pH 8.0) for 24 hours, after which the sample was separated into soluble and insoluble fractions by centrifugation. The insoluble pellet, containing fibrillar species, was denatured with guanidinium hydrochloride, resuspended in 20 mM MES (pH 6.0), and analyzed by HSQC NMR spectroscopy (Fig. 6*A*). The NMR spectra showed a significant decrease in peak intensities for the N- and C-terminal domain residues relative to the untreated fibril pellet, indicating that HtrA1 cleaves the NAC-flanking regions of α-Syn fibrils but not the β-sheet core formed by the NACore residues (Fig. 6*A*). Unlike the cleavage observed for HtrA1 and monomeric α-Syn (Fig. 4*A*,*B*), no cleavage at T92 was detected in α-Syn fibrils, likely due to its inaccessibility within the NACore of the fibril (7, 47). Additionally, SDS-PAGE confirmed that HtrA1 cleaves α-Syn fibrils in a time-dependent manner, with most cleavage occurring after 16 hours of incubation (Fig. S2). The selective cleavage of the N- and C-terminal domains of α-Syn fibrils suggests that HtrA1 activity is due to interactions with these domains on the fibril surface. To evaluate this hypothesis, α-Syn fibrils were incubated with HtrA1* in 10 mM PBS (pH 7.4) at 37°C, and the interaction was visualized using air AFM imaging (Fig. 6*B*-*D*). Amplitude images showed that HtrA1* colocalizes to several regions on the fibril surface, supporting the conclusion that HtrA1 interacts with the fuzzy coat of the N- and C-termini of α-Syn fibrils (Fig. 6*D*), a finding that suggests this binding may reduce the availability of aggregation-prone surfaces and thereby limits seeding activity.

**Figure 6:**
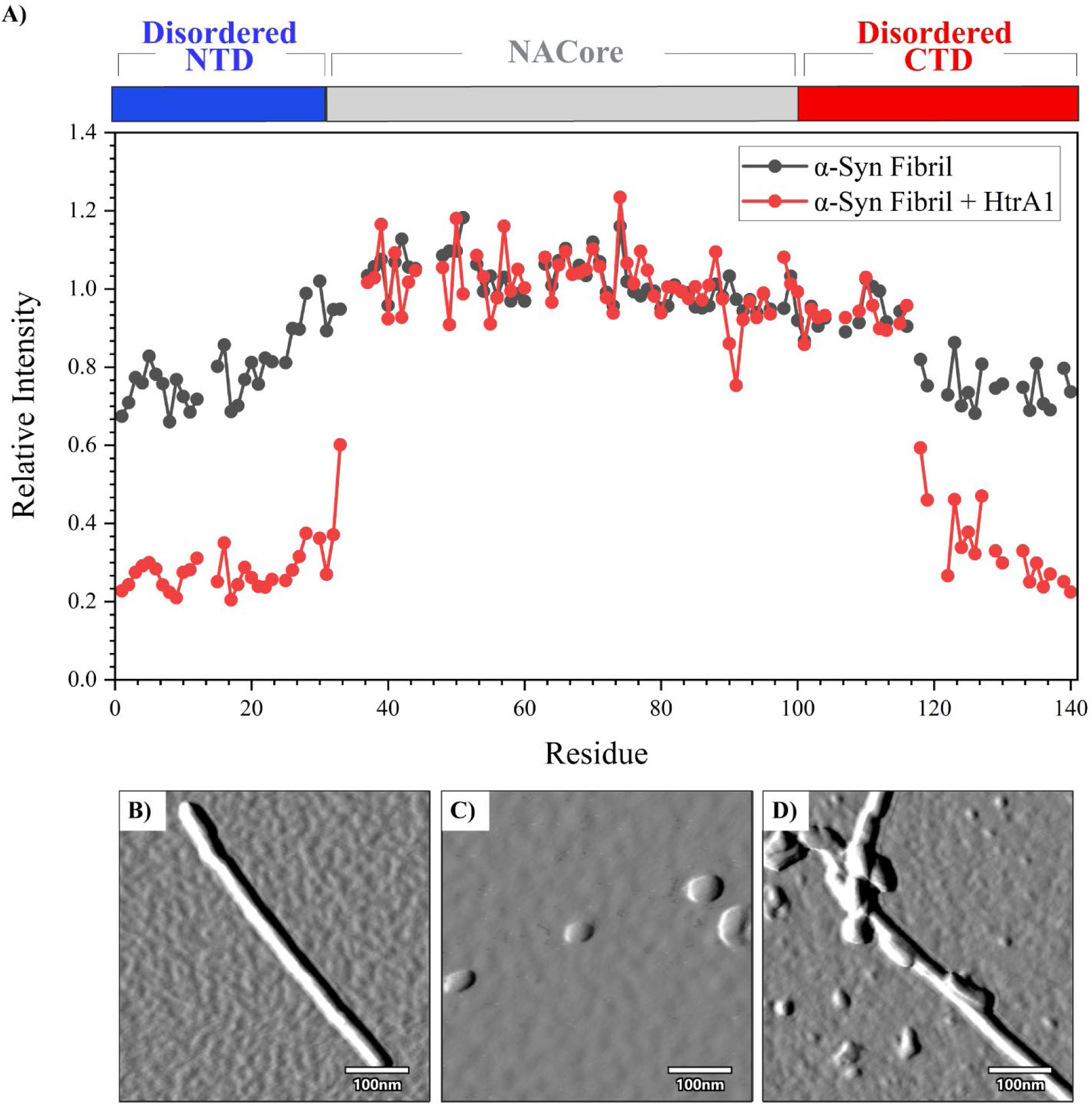
HtrA1 interacts with the intrinsically disordered N- and C-termini of α-Syn fibrils. (**A**) Peak intensity of ^15^N-^1^H HSQC spectrum peak intensities of ^15^N-labeled α-Syn fibrils in the absence (black) and presence (red) of HtrA1. Fibrils were incubated with (red) or without (black) 2.5 eq HtrA1 in 50 mM Tris-HCl (pH 8.0) for 24 hours. After, the soluble and insoluble portions of the sample were separated using centrifugation. The pellet was denatured in 8 M guanidinium hydrochloride, resuspended in NMR buffer (20 mM MES buffer, pH 6.0) and the NMR spectra were measured. The reduction in peak intensity at the N- and C-terminal regions of α-Syn fibrils upon HtrA1 addition indicates that HtrA1 selectively cleaves these disordered termini, disrupting fibril integrity. NMR data was acquired in NMR buffer with 10% D2O and at 15 °C, 700 MHz ^1^H Larmor frequency. (**B-D**) Amplitude air AFM images showing the colocalization of HtrA1* on the surface of α-Syn fibrils. In this setup, 5.0 μM α-Syn fibrils were incubated with 0.25 μM HtrA1* in 10 mM PBS (pH 7.4) (**D**), revealing that HtrA1* binds directly to the fibril surface. Imaging was performed at 25°C using AC240 tips (nominal resonance frequency: 70 kHz, spring constant: 2 N/m) on a Cypher ES AFM. Control images of α-Syn fibrils (**B**) and HtrA1* alone (**C**) confirmed that colocalization was specific to HtrA1* binding to fibrils (**D**).

## Discussion

This study identified the molecular mechanisms underlying the potent inhibitory effect of the molecular chaperone HtrA1 on α-Syn fibril formation and growth, focusing on key multi-domain interactions involved in various microscopic steps of the aggregation process. ThT aggregation inhibition experiments revealed that HtrA1*, a catalytically inactive variant, inhibits aggregation more effectively than the PDZ or Protease* domains, or even their combination, in both monomeric and seeded aggregation assays (Fig. 7). To understand this inhibition, the interactions of HtrA1*, PDZ, and Protease* with both monomeric and fibrillar α-Syn were examined. Our findings suggest that the inhibitory activity of HtrA1* arises from the ability of its individual domains to synergistically target different regions of α-Syn, allowing simultaneous targeting of both the monomeric and fibrillar forms.

**Figure 7:**
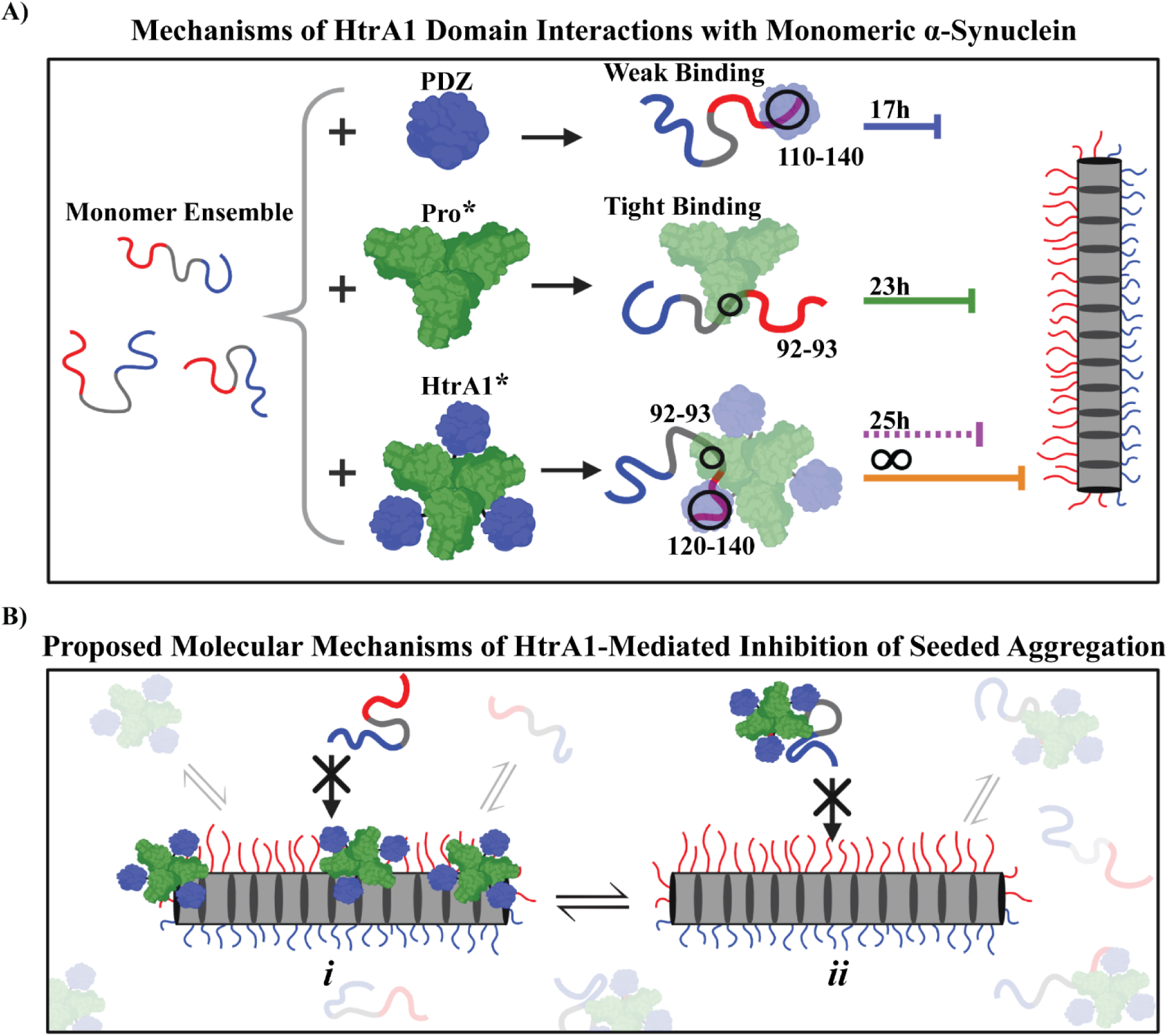
HtrA1* is a potent inhibitor of α-Syn aggregation by targeting multiple steps in the aggregation pathway. (**A**) Schematic illustrating HtrA1* and its domain-specific mechanisms for inhibiting fibril formation through interactions with monomeric α-Syn. HtrA1* interacts with α-Syn monomers by binding to residues 110–140 in the C-terminus and residues 92–93 in the NAC domain. The inhibitor line represents the half-time (T_1/2_) from the *de novo* ThT aggregation inhibition assay at a 1:1 α-Syn ratio, with colors indicating the following: PDZ (blue), Pro* (green), PDZ and Pro* combined (purple), and HtrA1* (orange). HtrA1* showed the strongest inhibitory effect followed by the combined domains, Pro* alone, and PDZ alone*. (**B**) Proposed molecular mechanisms of HtrA1-mediated inhibition of α-Syn seeded aggregation. In the presence of seeds, HtrA1* inhibits seeded aggregation through 2 mechanisms: (***i***) HtrA1* binds to the fibril surface, blocking available binding sites and preventing the recruitment of monomers for seeded aggregation; (***ii***) HtrA1*’s binding to the NAC and C-terminal regions of α-Syn monomers hinders their recruitment to the fibril’s fuzzy coat, thereby inhibiting seeded aggregation. The PDZ domain (dark blue) and Protease domain (green) mediate HtrA1*’s interactions with α-Syn. The α-Syn N-terminal domain is depicted in red, and the C-terminal domain in light blue.

We propose that HtrA1* disrupts α-Syn aggregation by binding synergistically to the C-terminus and NAC domain of α-Syn, with monomeric interactions facilitated through its PDZ and Protease* domains respectively (Fig. 7*A*). Solution-based NMR experiments and computational analyses reveal that the PDZ interacts with the C-terminal end of the α-Syn monomer, binding weakly to residues 136-140 and more broadly engaging residues 110-140 through its carboxylate-binding pocket and neighboring structures, respectively. Biophysical, computational, and NMR-based investigations reveal that HtrA1, *via* its Protease domain, targets and cleaves at a site towards the C-terminal end of the NAC domain of α-Syn monomer at T92-G93. Elegant experiments have demonstrated nanomolar binding of Protease* to the α-Syn monomer (25). Based on these findings, cleavage experiments, and the delayed half-time of aggregation observed in ThT fluorescence assays, Protease* is proposed to bind near residue 92 in the NAC domain of α-Syn to delay aggregation. HtrA1* achieves robust potent inhibition of α-Syn aggregation through synergistic binding, with PDZ targeting residues 110-140 in the C-terminal domain and Protease* engaging residues around 92-93 in the NAC domain.

Notably, the individual domains of HtrA1, with PDZ binding weakly at the C-terminus and Protease* binding strongly at the NAC region (25), are unable, whether individually or combined, to fully inhibit aggregation, as shown in ThT aggregation inhibition experiments (Fig. 7*A*). This limitation may arise because each domain targets only a single region of α-Syn, highlighting that inhibition may depend more on binding location than on binding affinity. HtrA1*, however, demonstrates the unique ability to simultaneously engage both the NAC and C-terminal regions of α-Syn monomers. This dual interaction more effectively inhibits aggregation through synergistic actions of the Protease* and PDZ domains, blocking residues 92–140 of α-Syn and preventing early aggregation events. Supporting this scenario, a peptide comprising α-Syn residues 1–92 (generated by Protease cleavage) does not independently aggregate. Strikingly, α-Syn treated with Protease* alone and the combined domains exhibited similar half-times of aggregation in ThT fluorescence, suggesting that PDZ’s weak binding adds minimal benefit when the domains are unlinked (Fig. 7*A*). However, when PDZ is linked to Protease* in HtrA1*, its weak binding becomes crucial in effectively inhibiting aggregation, emphasizing the critical role of synergistic interactions for potent inhibition. Conventionally designed inhibitors have typically focused on high binding affinity to a single region of α-Syn to inhibit aggregation (48–52). In contrast, our findings with HtrA1’s multidomain interactions suggest that targeting multiple domains, even with individually weak interactions, could offer a powerful new strategy for inhibitor design. This multi-pronged mechanism is exemplified by natural inhibitors like dopamine, which binds residues 125–129 in the C-terminus and interacts electrostatically with residue 83 in the NAC region, and beta synuclein (β-Syn), which transiently stabilizes α-synuclein by interacting with multiple sites to prevent aggregation. (53, 54).

HtrA1* effectively suppresses α-Syn seeded aggregation by targeting different steps in the process (Fig. 7*B*). Studies have suggested that fibril growth during seeded aggregation occurs through recruitment of the monomer N-terminal region to the fibril ‘fuzzy coat’ (55), in particular to the disordered C-terminus of the fibril (32, 38, 39). We propose that HtrA1* disrupts monomer recruitment by binding both to the fibril surface (Fig. 7B, *i*) and to α-Syn monomers (Fig. 7B, *ii*). Inhibition of fibril surface binding is supported by HtrA1* visualization on the fibril surface, NMR evidence suggesting PDZ binding at the C-terminus, and cleavage assays demonstrating that HtrA1 broadly targets the fibril’s intrinsically disordered regions (IDRs).These interactions compete with monomer recruitment by blocking available binding sites on the fibril surface (Fig. 7B, *i*). Additionally, HtrA1* binding to α-Syn monomers may obstruct their incorporation into the fibril, further inhibiting seeded aggregation (Fig. 7B, *ii*). Collectively, the ability of HtrA1* to disrupt multiple steps in seeded aggregation positions it as a promising model for therapeutic design, highlighting the importance of targeting α-Syn’s IDRs to develop effective amyloid aggregation inhibitors.

Previous elegant studies have reported mixed findings on the role of the HtrA1 PDZ domain in mechanistic interactions, with some studies demonstrating its necessity for functionality (26, 31, 35) while other reports suggest it is non-essential in the context of inhibition (25). Our data supports the view that PDZ inhibits both unseeded and seeded α-Syn aggregation in a dose-dependent manner and that potent inhibition by HtrA1* arises due to cooperative interactions between PDZ and Protease*. While PDZ is the least inhibitory of the three constructs, it still inhibits unseeded α-Syn aggregation in a dose-dependent manner through its weak binding to the C-terminal end of α-Syn. Notably, the PDZ domain alone does not efficiently inhibit seeded aggregation at substoichiometric concentrations due to its weak binding, which may allow it to be outcompeted by unbound α-Syn monomers; however, it shows inhibition at higher concentrations. In the context of HtrA1* when PDZ is linked to the trimeric protease, oligomerization may effectively enhance its weak affinity due to avidity effects. PDZ also enhances the inhibition of seeded aggregation by HtrA1*, as shown by limited inhibition with Protease* at low concentrations, while HtrA1* achieves full inhibition. These results suggest that while PDZ monomer alone leads to weak inhibition, it is crucial for achieving complete inhibition through its synergistic interactions with the trimeric Protease*.

Over the past decade, substantial research has focused on the aggregation-prone core regions of α-Syn fibrils to understand the mechanisms driving α-Syn aggregation and to develop therapeutics for neurodegenerative diseases (10, 12, 56–58). Sophisticated strategies include developing small molecule inhibitors that target the NAC domain to hinder secondary nucleation (12), designing mini proteins that bind to fibril ends to block elongation (13), and utilizing chaperones like Hsp27 to bind to fibril ends and prevent further growth suggesting chaperones as promising inhibitors (59). Although the NAC domain stabilizes the core of α-Syn fibrils, recent studies indicate that the disordered flanking regions play essential roles in nucleation and elongation (10, 32, 38, 60). Our findings elucidate HtrA1’s inhibitory mechanisms, highlighting the therapeutic potential of targeting both the NAC and its flanking regions in α-Syn as a potent strategy for mitigating amyloid formation.

## Materials and Methods

### Protein expression and purification

Expression of N-terminally acetylated human α-Syn was performed *via* co-expression with pNatB plasmid (Addgene #53613) in *Escherichia coli* (*E. coli*) BL21(DE3) cells (New England Biolabs, MA) and protein purification was performed as described previously (61). Uniformly ^15^N or ^13^C,^15^N isotopically labeled acetylated α-Syn for NMR experiments was expressed in M9 minimal media supplemented with ^15^N-ammonium chloride (Millipore Sigma, St. Louis, MO) or ^13^C D-glucose (Cambridge Isotopes Laboratories, Tewksbury, MA) as the sole nitrogen and carbon sources, respectively. Protein molecular weight and purity were assessed by ESI-MS and the purified protein was stored at −20°C as a lyophilized powder until use.

Expression of non-acetylated human α-Syn occurred *via* transformation into *E. coli* BL21(DE3) cells (New England Biolabs, MA) and plated onto agar plates containing the antibiotic ampicillin (50 ng/mL). 50 mL starter cultures were inoculated using one colony each and grown in LB media at 37°C, 160 rpm overnight (Scientific Innova 44 Incubator Shaker, New Brunswick, NJ). 1L LB cultures were inoculated using 10 mL starter cultures and were allowed to grow at 37°C, 180 rpm until an O.D of 0.6 was reached. At this point, the cultures were induced with 1 mM IPTG and grown at 20°C, 180 rpm for 16-18 hours. After overexpression, cultures were pelleted down and stored at −80 °C until use.

Non-acetylated human α-Syn was purified by resuspending the bacterial pellets in 10 mM PBS (pH 7.4). The protein solutions were lysed *via* sonication (Sonicator Dismembrator Model 500, Fisher Scientific, MA) at 30% amplitude for 8 cycles of 15 seconds on 30 seconds off. Solutions were kept on ice to prevent degradation. Following lysis, samples were boiled for 20 minutes and centrifuged at 20,000 rpm for 45 minutes (Avanti J-26S XPI Centrifuge, Beckman Coulter, CA) to remove cellular debris. The supernatants were collected and treated with streptomycin sulfate (10 mg/mL) to remove nucleic acids. The solutions were mixed at 4°C for 20 minutes and centrifuged as previously described. The protein was precipitated by adding ammonium sulfate (0.361 g/mL) and mixed at 4°C for 1 hour after which the solutions were centrifuged once more to collect the final protein pellets. The pellets were resuspended in Buffer A (32 mM Tris, pH 7.8) and prepared for fast protein liquid chromatography (FPLC) by filtration using a 0.22 µm syringe filter. The filtered protein solutions were injected onto a HiTrap Q HP anion exchange column (Cytiva AKTA, MA) and the purified protein was eluted at 50% Buffer B (32 mM Tris, 500 mM NaCl, pH 7.8) using an AKTA Pure FPLC (Cytiva AKTA, MA). The purified protein was flash frozen using LN_2_ and stored at −80 °C until use.

Human HtrA1 without an N-terminal domain with a N-terminal Strep-II tag was provided by Prof. M. Ehrmann in expression vector pET21d. The PDZ and Protease domain of HtrA1 were amplified using primers 5’-ATCACCAAGAAGAAGTATATTG-3’ and 5’-GGATCCTTTTTCGAACTGC-3’ as well as 5’-TAGCTCGAGCACCACCAC-3’ and 5’-TTTGGCCTGTCGGTCATG-3’with a NEB Q5® Site-Directed Mutagenesis Kit (New England Biolabs, MA). The mutation of S328A of HtrA1 and its Protease domain were created using primers 5’-CTATGGAAACgcgGGAGGCCCGT-3’ and 5’-TTGATGATGGCGTCGGTCTG-3’ with a NEB Q5® Site-Directed Mutagenesis Kit (New England Biolabs, MA). The PCR protocol was as followed: 95°C for 5 minutes, 35 cycles of 30 seconds at 95°C, 30 seconds at 52 °C, and 60 seconds at 72°C, followed by 72°C for 5 minutes. Plasmids were transformed into *E. coli* BL21(DE3) cells (New England Biolabs, MA) for expression in LB media or M9 minimal media containing either ^15^N-ammonium chloride (Millipore Sigma, St. Louis, MO) or ^13^C D-glucose (Cambridge Isotopes Laboratories, Tewksbury, MA) for isotopic labeling. 1L LB or M9 minimal media cultures were inoculated using 10 mL starter cultures and were allowed to grow at 37°C, 80 rpm until an O.D of 0.6 was reached. At this point, the cultures were induced with 1 mM IPTG and grown at 20°C, 180 rpm for 16-18 hours. After overexpression, cultures were pelleted down and stored at −80 °C until use.

HtrA1 and its variants were purified by resuspending the bacterial pellets in 50 mM Tris, 300 mM NaCl buffer (pH 7.5). The protein solutions were lysed *via* sonication (Sonicator Dismembrator Model 500; Fisher Scientific, MA) at 30% amplitude for 8 cycles of 15 seconds on 30 seconds off. Solutions were kept on ice to prevent degradation. Following lysis, samples were centrifuged at 20,000 rpm for 45 minutes (Avanti J-26S XPI Centrifuge, Beckman Coulter, CA) to remove cellular debris. The cell lysate was filtered using a 0.22 µm syringe filter and loaded onto a StrepTrap HP column (Cytiva AKTA, MA). The column was washed with 50 mM Tris, 300 mM NaCl buffer (pH 7.5) and the protein was eluted off the column with elution buffer (50 mM Tris, 300 mM NaCl buffer, 2.5 mM desthiobiotin (pH 7.5)) using an AKTA Pure FPLC (Cytiva AKTA, MA). The purified protein was flash frozen using LN_2_ and stored at −80°C until use.

### Fibril sample preparation

Lyophilized acetylated α-Syn was dissolved in 10 mM PBS (pH 7.4), and large aggregates were removed by using a 100 kDa centrifuge filter (Millipore Sigma, St. Louis, MO). The dissolved protein was concentrated using a 3 kDa centrifuge filter (Millipore Sigma, St. Louis, MO) to 70 uM. 100 uL of each sample mixture was loaded into 96-well clear bottom plates (Corning, Corning, NY) with a single Teflon bead (3 mm, Saint-Gobain N.A., Malvern PA). The plates were sealed with Axygen sealing tape (Corning, Corning, NY) and shaken at 37°C 600 rpm in a POLAR Star Omega plate reader (BMG Labtech, Cary, NC). Fibrils samples were collected by centrifugation at 20,000g for 40 minutes (Microcentrifuge 5430, Eppendorf, MA). After removing the supernatant, the fibril pellets were washed through multiple rounds of re-suspension in 10 mM PBS (pH 7.4) and centrifuged at 20,000g for 40 minutes to remove residual soluble and non-fibrillar components. The fibril pellets were stored at room temperature until use.

### ThT fluorescence assays

Lyophilized acetylated α-Syn was dissolved in 10 mM PBS (pH 7.4), and large aggregates were removed by using a 100 kDa centrifuge filter (Millipore Sigma, St. Louis, MO). The dissolved protein was concentrated using a 3 kDa centrifuge filter (Millipore Sigma, St. Louis, MO). HtrA1 variants (PDZ, inactive Protease (Pro*), and inactive HtrA1 (HtrA1*)) were buffer exchanged and concentrated in 10 mM PBS (pH 7.4) using a 10 kDa filter (Millipore Sigma, St. Louis, MO). For *de novo* aggregation ThT assays, α-Syn monomer was diluted to 50 uM, mixed with 20 μM ThT (Acros Organics, Pittsburgh, PA), and loaded into 96-well plates (Corning, Corning, NY). HtrA1 variants were individually added at various concentrations. A single Teflon bead (3 mm; Saint-Gobain N.A., Malvern, PA) was added to each well, and plates were sealed with Axygen sealing tape (Corning, Corning, NY). Plates were shaken at 37°C 600 rpm for 60 hours, and the increase in ThT fluorescence intensity at 480 nm was measured every 33 minutes by a POLAR Star Omega plate reader (BMG Labtech, Cary, NC). At least three replicates of the ThT assays were recorded for each sample. Fluorescence traces were averaged, and standard deviation of these traces are presented.

For seeded aggregation ThT assays, acetylated α-Syn monomer was diluted to 50 uM, mixed with 2.5 uM sonicated fibrils (seeds), and 20 μM ThT (Acros Organics, Pittsburgh, PA). Samples were loaded into 96-well plates (Corning, Corning, NY). HtrA1 variants were individually added at various concentrations. Plates were sealed with Axygen sealing tape (Corning, Corning, NY) and incubated at 37°C under quiescent conditions for 60 hours, and the increase in ThT fluorescence intensity at 480 nm was measured every 33 minutes by a POLAR Star Omega plate reader (BMG Labtech, Cary, NC). At least three replicates of the ThT assays were recorded for each sample. Fluorescence traces were averaged, and standard deviation of these traces are presented.

### Proteolysis experiments

For the SDS-PAGE monomeric proteolysis experiments, acetylated α-Syn monomer was incubated with active HtrA1 and active Protease at a 5:1 ratio in 50 mM Tris-HCl (pH 8.0) at 37°C for various time points (0, 15 minutes, 30 minutes, 1 hour, 2 hours, 4 hours, and overnight). After proteolysis, the reaction products were heated in 1X Laemmli Buffer (Bio-Rad, Hercules, CA) at 90°C for 5 minutes (Thermomixer C, Eppendorf, MA). The samples were loaded onto a 4-20% Mini-PROTEAN precast protein gel (Bio-Rad, Hercules, CA) and ran on a Mini-PROTEAN Tetra Vertical Electrophoresis Cell (Bio-Rad, Hercules, CA). After, gels were fixed and stained using Brilliant Blue Coomassie (Fisher Scientific, Hanover Park, IL).

To identify the proteolysis products, acetylated α-Syn monomer was incubated with active HtrA1 and active Protease at a 5:1 and 2:1 ratio, respectively, in 50 mM Tris-HCl (pH 8.0) at 37°C for various timepoints. The proteolysis reactions were stopped by heating the sample at 95 °C for 10 minutes. The denatured HtrA1 was removed by centrifugation at 20,000 rpm for 10 minutes (Microcentrifuge 5430, Eppendorf, MA). The supernatant was passed through PD-10 Desalting Columns (Millipore Sigma, St. Louis, MO) to remove excess salts. The samples were dialyzed using a 3.5 kDa MWCO tubing overnight against 50 mM ammonium acetate with 0.1% formic acid. The sample was diluted to a final protein concentration of 10 μM for electrospray ionization mass spectrometry (SYNAPT G2-SI qTOF, Waters Corp, MA).

For the SDS-PAGE fibrillar proteolysis experiments, sonicated α-Syn fibrils were incubated with active HtrA1 at a 5:1 ratio in 50 mM Tris-HCl (pH 8.0) at 37°C for various time points (0, 30 minutes, 1 hour, 2 hours, 3 hours, 4 hours, and 16 hours). After proteolysis, the reaction products were heated in 1X Laemmli Buffer (Bio-Rad, Hercules, CA) at 95°C for 10 minutes (Thermomixer C, Eppendorf, MA). Before loading onto a protein gel, the samples were denatured with 4 M guanidinium hydrochloride. The samples were loaded onto a 4-20% Mini-PROTEAN precast protein gel (Bio-Rad, Hercules, CA) and ran on a Mini-PROTEAN Tetra Vertical Electrophoresis Cell (Bio-Rad, Hercules, CA). After, gels were fixed and stained using Brilliant Blue Coomassie (Fisher Scientific, Hanover Park, IL).

### Thermomixer experiment and TEM imaging

To observe how HtrA1* affects fibril formation in the absence of ThT dye, 50 μM non-acetylated α-Syn was incubated in the absence and presence of 50 μM HtrA1* in 10 mM PBS (pH 7.4) at 37°C 900 rpm for one week in a thermomixer (Thermomixer C, Eppendorf, MA). After one week of fibrilization, the solutions were spun down at 15000 rpm for 2 hours (Microcentrifuge 5430, Eppendorf, MA) and the pellets containing the insoluble species were collected. The pellets were resuspended in 10 mM PBS (pH 7.4) and imaged using TEM. For TEM sample preparation, 3.5 µL of each sample was added to a CF mesh 200-Cu Carbon coated grid (Electron Microscopy Sciences) and incubated at room temperature for 90 seconds. Excess liquid was blotted off using filter paper. The grids were negatively stained for one minute using 3.5 µL of 3% uranyl acetate (Electron Microscopy Sciences). Excess liquid was blotted off using filter paper and the grids were dried overnight before imaging. TEM images were taken using a JEOL JEM2010 TEM (JEOL, Japan) at 200.0 kV.

### PDZ pulldown experiment

50 or 100 uM PDZ was incubated with 50 uM acetylated α-Syn fibril in 10 mM PBS (pH 7.4) at 37°C 300 rpm for 24 hours (Thermomixer C, Eppendorf, MA). 50 uM PDZ only and 50 uM acetylated α-Syn fibril only samples with the same treatment were used as controls. The fibril concentrations reported are in monomer equivalents as determined by a BCA assay (Pierce™ BCA Protein Assay Kit, Thermo Fisher Scientific, MA). After incubation, all of the samples were pelleted at 20,000 rpm for 1 hour (Microcentrifuge 5430, Eppendorf, MA). Before running on SDS-PAGE, the pellets were denatured using 8 M urea. The dissociated pellets were buffer exchanged to 10 mM PBS (pH 7.4) using a 3 kDa centrifuge filter (Millipore Sigma, St. Louis, MO). The samples were loaded onto a 4-20% Mini-PROTEAN precast protein gel (Bio-Rad, Hercules, CA) and ran on a Mini-PROTEAN Tetra Vertical Electrophoresis Cell (Bio-Rad, Hercules, CA). After, gels were fixed and stained using Brilliant Blue Coomassie (Fisher Scientific, Hanover Park, IL).

### Air AFM imaging

Air AFM imaging was used to show the colocalization of HtrA1* to the fibril surface. Briefly, 5.0 μM of acetylated α-Syn fibrils were incubated with 0.25 μM of HtrA1* in 10 mM PBS (pH 7.4). Approximately 50 µL of the fibril-HtrA1* sample was placed onto freshly cleaved mica and allowed to bind to the surface for 5 minutes. After, the mica surface was washed with 2 mL of ultrapure water (Millipore Sigma, St. Louis, MO) and the sample was allowed to dry at room temperature for one hour in a laminar flow hood. The sample was then placed into the AFM for imaging (Cypher ES AFM, Asylum Research, Oxford Instruments, United Kingdom). Imaging was performed at 25°C with AC240 tips (nominal resonance frequency of 70 kH, nominal spring constant of 2 N/m). Control imaging was performed on the acetylated fibrils and HtrA1* as well.

### Timelapse liquid AFM imaging

For the timelapse liquid AFM imaging sample preparation, 50 uL of 5 µM sonicated non-acetylated α-Syn fibrils (seeds) were placed onto a freshly cleaved mica surface and allowed to bind for 5 minutes. After, the mica surface was washed with 2 mL of ultrapure water (Millipore Sigma, St. Louis, MO) and 1 mL of 10 mM PBS (pH 7.4) was added to the surface. The sample was placed into the AFM to confirm the deposition of seeds on the mica surface. After confirmation, the liquid on the surface was exchanged for 100 uL of 25 µM non-acetylated α-Syn monomer in 10 mM PBS (pH 7.4) or in a mixture containing monomer and 1 µM Pro* or HtrA1* in 10 mM PBS (pH 7.4). After which the samples were imaged to monitor the growth of seeds.

All AFM timelapse experiments were performed on a Cypher ES AFM (Asylum Research, Oxford Instruments, United Kingdom) using PNP-DB cantilevers (NanoAndMore, CA) with a nominal spring constant of k = 0.48 N/m and a nominal resonance frequency of r < 10 nm. Imaging was performed at 25°C in tapping mode which utilized blueDrive (Asylum Research, Oxford Instruments, United Kingdom) photothermal excitation to aid in producing high quality images in liquid and for determining the resonance frequency of individual tips. To monitor the growth of the seeds, an area of the mica was selected which contained seeds and imaged continuously for several hours. After imaging, individual AFM images were stitched together to form videos illustrating the growth.

### NMR sample preparation and experiments

For the preparation of the α-Syn NMR experiments, lyophilized ^15^N-labelled acetylated α-Syn monomer powder was dissolved in NMR buffer (20 mM MES, 100 mM NaCl, pH 6.0). The protein was filtered through a 100 kDa centrifuge filter (Millipore Sigma, St. Louis, MO) to remove higher-order oligomers, and then concentrated using a 3 kDa centrifuge filter (Millipore Sigma, St. Louis, MO). Samples were diluted to a final protein concentration of 200 μM with 10% D_2_O added. NMR spectra were recorded on an 800 MHz Varian Inova spectrometer. Spectra were processed using NMRPipe. For the addition of PDZ to α-Syn, 1 eq of nonlabelled PDZ was buffer exchanged to the NMR buffer using a 3 kDa centrifuge filter (Millipore Sigma, St. Louis, MO) and directly added to the monomer.

For the ^15^N-^1^H HSQC of proteolysis of α-Syn fibrils by HtrA1, ^15^N-labelled α-Syn fibril were incubated with buffer or 2.5 eq of HtrA1 in 50 mM Tris-HCl (pH 8.0) for 24 hours. The soluble and insoluble portions were separated using centrifugation. The insoluble portion was denatured using 8 M guanidinium hydrochloride and resuspended in NMR buffer. NMR spectra were measured at 800 MHz with 64 scans.

For the preparation of PDZ domain NMR experiments, ^15^N-labelled PDZ was buffer exchanged to 20 mM MES, 100 mM NaCl (pH 6.0) using a 3 kDa centrifuge filter (Millipore Sigma, St. Louis, MO). Samples were diluted to a final protein concentration of 200 μM with 10% D_2_O added. NMR spectra were recorded on an 800 MHz Varian Inova or a 700 MHz Bruker Avance III spectrometer. Spectra were processed using NMRPipe. For the addition of α-Syn monomer, 1 eq of nonlabelled α-Syn monomer was buffered exchange to the NMR buffer using a 3 kDa centrifuge filter (Millipore Sigma, St. Louis, MO) and directly added to the PDZ. For the addition of α-Syn fibril, 0.25 or 0.35 monomer eq of nonlabelled α-Syn fibrils in NMR buffer was directly added to the ^15^N-labelled PDZ.

### Computational methods

#### Model preparation

##### Protease

Construction of the protease model began with the PDB structure 3NZI (62). The boron-modified peptide in the original model was replaced with an unmodified peptide, generated with Rosetta to match the sequence and backbone dihedrals, superimposed over the original. Geometric EnzDes constraints were developed for the catalytic site and oxyanion hole (residues H220, D250, G326, S328, and P1) to preserve near-attack geometry, and the whole structure was minimized using Rosetta’s FastRelax protocol with coordinate and catalytic constraints, using the best scoring model of 100 trajectories. Models with elongated substrates were produced by generating a peptide with Rosetta to match the α-Syn sequence in the region to be modeled, setting the backbone dihedrals of the P5-P1 residues to match the original 3NZI and all other residues at −180° for all backbone dihedrals, and replacing the original peptide with the elongated one, superimposed in the P5-P1 region before applying the sampling protocol.

##### PDZ

Construction of the PDZ model began with the PDB structure 2JOA (35). All NMR ensemble members of the original model were minimized in Rosetta similarly to the protease, with 20 trajectories per member. For subsequent modeling, the best-scoring models from the four best-scoring states were used (2, 11, 16, and 18). Models with extended substrate peptides were generated in a similar fashion to those of the protease.

### Identifying protease site specificity

#### Protease ensemble generation

To assess the correlation of cleavage site preferences with protease binding energy, we modeled all α-Syn subframes bound in the protease active site. We used protease models including substrates starting at residues P6 or P5 and extending to all residues P1’-P6’ (a total of 12 substrate frames with frame lengths between 6 and 12) and P4 to P1’ were used in frame scanning. For each possible α-Syn frame (for a frame length of 6, frames would include α-Syn residues 1-6, 2-7, 3-8, etc.), the substrate sequence in the model was changed to the appropriate α-Syn sequence using Rosetta’s PackRotamersMover. We used Rosetta’s FlexPepDocking protocol for structural sampling of the peptide, with 100 trajectories per frame. We analyzed the ten lowest-energy models for each.

#### Site-based discriminator

Our discriminator values were determined similarly to previous work (46), using three components in a weighted sum: protease interfacial residue total scores (weight 1), substrate residue total scores (weight 1), and total constraint scores (weight 3.5). The protease interfacial residues were those with C_α_ within 8Å of the substrate. Constraint scores reflect the energetic penalty necessary to maintain a given substrate bound in catalytic conformation during structural sampling. Sites were ranked by the single best discriminator value determined among the ten lowest-energy structures.

### PDZ ensemble generation

#### PDZ ensemble generation

To assess the extent to which the NMR shifts identified in the C-terminal region of α-Syn and in the PDZ are attributable to binding energy (as opposed to other factors), we modeled an ensemble of PDZ-bound α-Syn C-terminal peptides. Since the NMR indicated that PDZ residues distal to the α-Syn C-terminus binding site, and α-Syn residues as far as 30 away from the C-terminus were affected by binding, we modeled a C-terminal section comprising 30 residues, bound with the PDZ. Initial peptide backbone angles were selected randomly from the unbound α-Syn structural ensemble, computationally generated by Ferrie and Petersson (63) and then structures were further refined with Rosetta’s FlexPepDocking protocol. We generated 12500 models and analyzed the 500 lowest-scoring structures. We determined the interfacial Rosetta energies of each model on a per-residue basis. The score for each PDZ residue excludes pairwise interaction energies with all other PDZ residues and all one-body energies.

## Acknowledgments

This work was funded and supported by the NSF (CHE2226816 to S.D.K.) and the NIH (GM136431 to J.B.; NIH T32 Training in Translating Neuroscience to Therapies Grant T32NS115700 to P.C.; NIH T32 Postdoctoral Training in Translational Research in Regenerative Medicine Grant T32EB005583 to J.R.). We would like to thank Dr. Michael Ehrmann of the University of Duisburg-Essen for providing us with the plasmid for HtrA1.

## Author Contributions

P.C., B.W., J.H.L., X.Y., J.R., S.D.K., and J.B. designed research; P.C., B.W., J.H.L., X.Y., J.R., S.D.K., and J.B. performed research; P.C., B.W., J.H.L., X.Y., J.R., S.D.K., and J.B. analyzed data; and P.C., B.W., J.H.L., S.D.K., and J.B. wrote the paper.

