## Supplemental Information for "Synergistic Multi-Pronged Interactions Mediate the Effective Inhibition of Alpha-Synuclein Aggregation by the Chaperone HtrA1"

**This PDF file includes:**

Figures S1 and S2
Legends for Movies S1 to S3

**Other supporting materials for this manuscript include the following:**

Movies S1 to S3

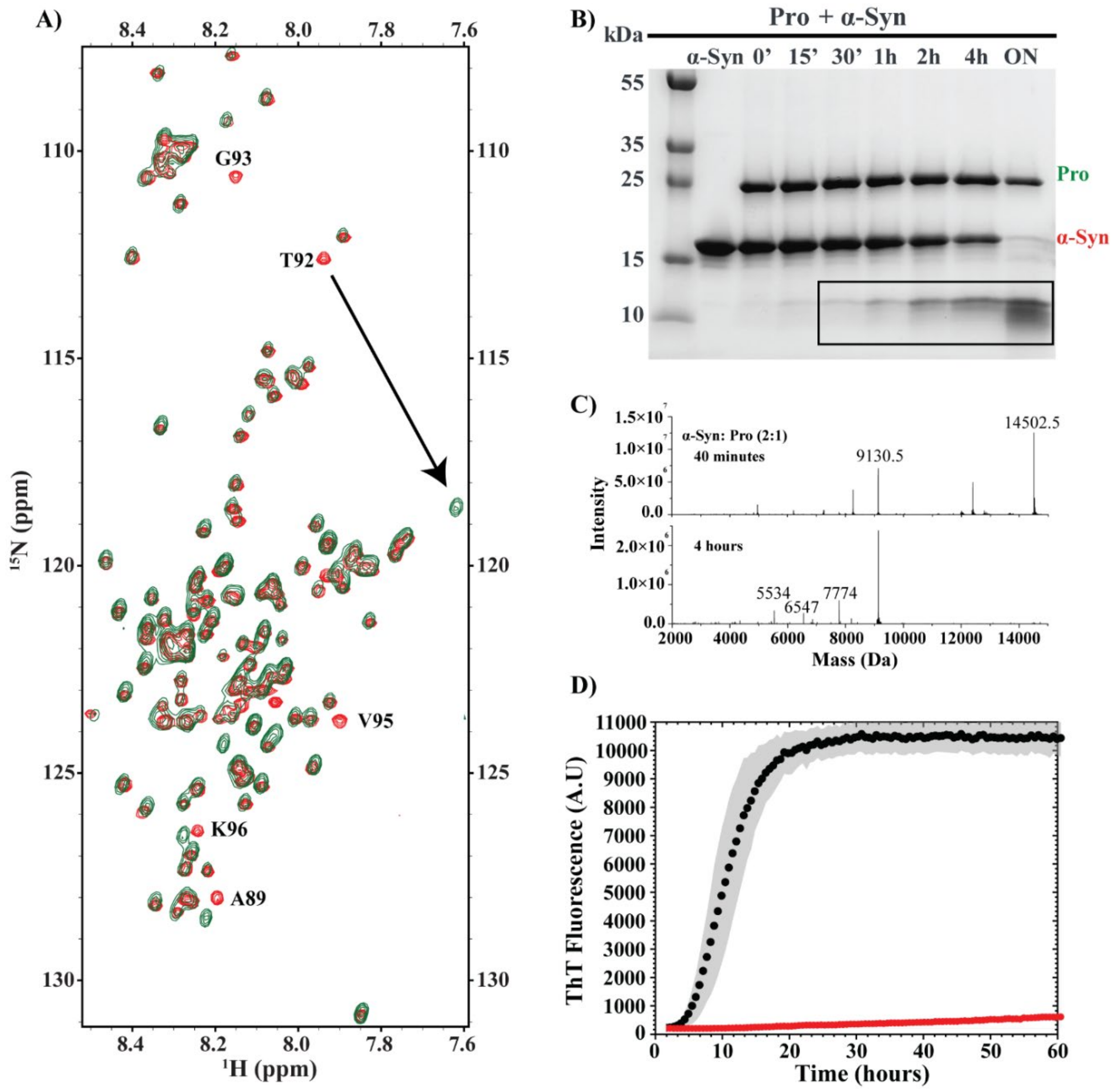


**Fig. S1.** HtrA1-mediated proteolysis of α-Syn *via* its Protease domain. (**A**) NMR spectra identifying the Protease domain's cleavage site on α-Syn. Overlay of $^{15}$N-$^{1}$H HSQC spectra of $^{15}$N labeled α-Syn monomer in the absence (red) and presence of Protease (green). Chemical shift perturbation of residue T92 and broadening of G93 are highlighted. (**B**) Proteolysis of α-Syn by Protease (Pro) visualized by SDS-PAGE. α-Syn monomer was incubated with Pro at a 5:1 (α-Syn: Pro) ratio over various timepoints in 50 mM Tris-HCl (pH 8.0) at 37°C. Cleavage fragments appeared within 30 minutes, with most monomer cleaved after overnight incubation. Bands corresponding to Pro (green) and α-Syn (red) are labeled, with cleaved α-Syn fragments highlighted by a black box. (**C**) Mass spectrometry analysis of Pro-mediated α-Syn cleavage. α-Syn monomer was incubated with Pro at a 2:1

(α-Syn: Pro) ratio in 50 mM Tris-HCl (pH 8.0) at 37°C for 40 minutes and 4 hours, yielding a main peak at 9130.5 Da, corresponding to the 1–92 α-Syn fragment. Incubation for 4 hours revealed additional peaks, indicating further cleavage beyond the T92–G93 site. (**D**) *De novo* fibril formation monitored by Thioflavin T (ThT) fluorescence. 50 μM of α-Syn monomer (black) and 50 μM of the 1–92 α-Syn peptide fragment (red) was incubated in 10 mM PBS (pH 7.4) at 37°C with constant agitation using a Teflon bead for 60 hours. No aggregation was observed for the peptide fragment. Together, these data highlight the specificity and functional consequences of Pro-mediated cleavage.

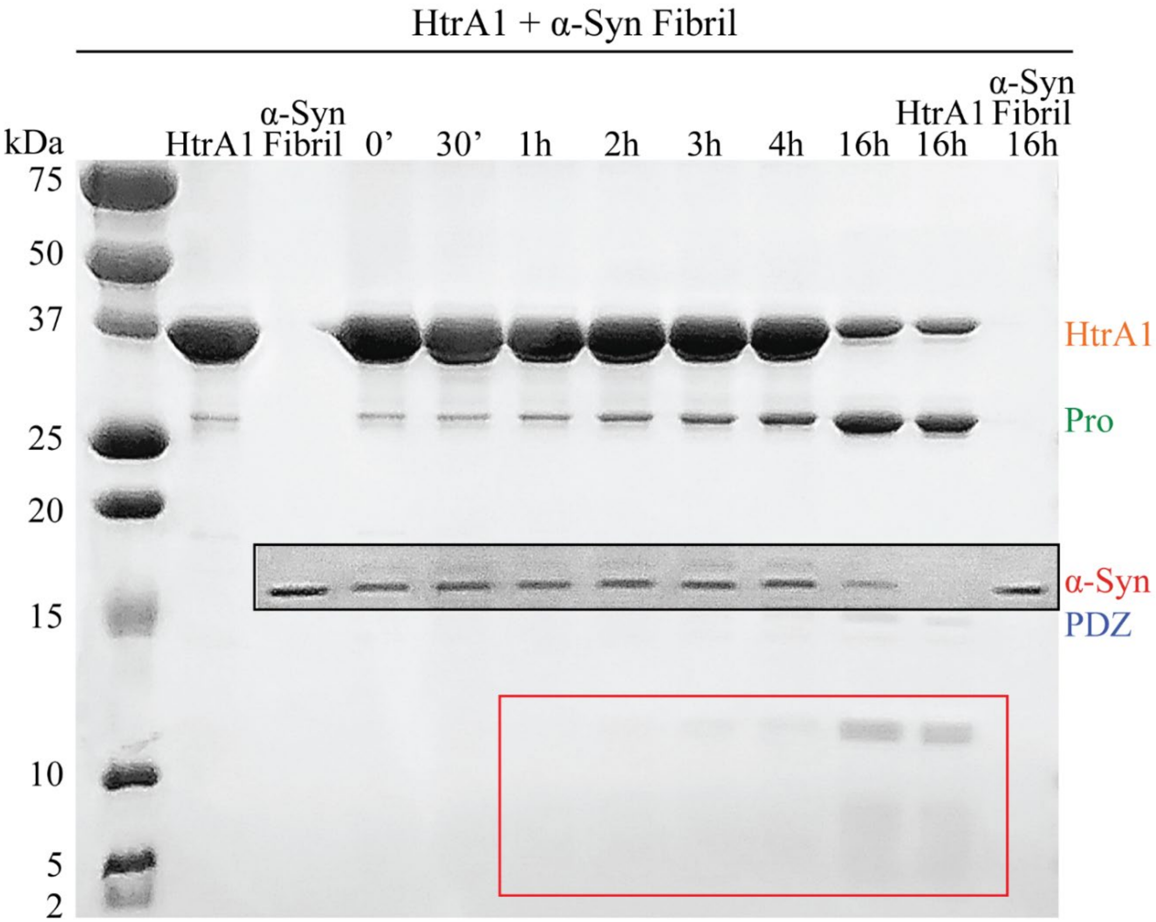


**Fig. S2.** Time-dependent proteolysis of α-Syn fibrils by HtrA1 visualized by SDS-PAGE α-Syn fibrils were incubated with HtrA1 at a 5:1 (α-Syn:HtrA1) ratio in 50 mM Tris-HCl (pH 8.0) at 37°C for various timepoints. Faint cleavage fragments of α-Syn were detected at 3 hours, with a majority of fibril cleavage occurring after 16 hours. Bands corresponding to HtrA1 (orange), Pro (green), α-Syn (red), and PDZ (blue) are labeled accordingly. he time-dependent decrease in fibril band intensity is marked by a black box, while cleaved α-Syn fragments are marked with a red box.

**Movie S1-3 (separate files).** AFM visualization of α-Syn fibril growth in the presence and absence of Protease* and HtrA1*. Seeded fibril growth was examined by placing 5 μM sonicated α-Syn fibrils (seeds) on mica, followed by adding 25 μM α-Syn monomer (Movie S1), 25 μM α-Syn with 1 μM Pro* (Movie S2), or 25 μM α-Syn with 1 μM HtrA1* (Movie S3) in 10 mM PBS (pH 7.4). Growth was imaged *via* AFM on a Cypher ES AFM (Asylum Research, Oxford Instruments) with PNP-DB cantilevers (k = 0.48 N/m, r < 10 nm). Imaging occurred at 25°C in tapping mode, with BlueDrive photothermal excitation ensuring high-quality liquid-phase images. These videos demonstrate that α-Syn monomer alone promotes rapid fibril growth, whereas Pro* partially inhibits fibril growth, and HtrA1* effectively prevents fibril formation.
